# Homeostatic, phagocytic, *NRF2/Hmox1*, *Apoc1* and chemokine microglia transcriptional programmes in naïve mouse cerebral cortex from embryo to adulthood

**DOI:** 10.64898/2026.08.20.746090

**Authors:** Varsini Sakthivadivel Ramasamy, Maide Ozen

## Abstract

Microglia, the resident immune cells of the central nervous system, undergo dynamic transcriptional remodeling across embryonic and postnatal development. However, the precise transcriptional programmes governing these transitions, and the role of oxidative stress pathways such as NRF2/Hmox1 in shaping microglial maturation, remain incompletely understood. Here, we characterized the transcriptional landscape of mouse microglial development using pseudobulk RNA-sequencing data, spanning five developmental stages, from embryonic day 17 to postnatal day 60. We identified four distinct transcriptional programmes (homeostatic, phagocytic, NRF2/Hmox1 oxidative stress-responsive, and Apoc1-associated) whose relative activities shift coordinately across development. Early developmental microglia were dominated by phagocytic and NRF2/Hmox1-associated gene expression, while mature microglia progressively acquired a homeostatic transcriptional identity marked by Tmem119 and P2ry12. Pseudotime trajectory analysis confirmed a continuous developmental axis along which the phagocytic programme declined, homeostatic programme increased, and NRF2/Hmox1 activity peaked at intermediate stages. Differential expression analysis distinguished Tmem119^+^ homeostatic microglia from Tmem119^-^ populations, and early developmental from mature microglial states. Additionally, chemokine receptor expression, including Cxcr4 at early timepoints, suggested a role for chemokine signaling in microglial migration and tissue integration during brain development. Collectively, these findings support a model in which microglial maturation proceeds along a transitional regulatory role during brain development.

## 2. Introduction

Microglia are the resident immune cells of the central nervous system and play a fundamental role in brain development[1]. Beyond their classical immune surveillance functions, microglia actively contribute to synaptic pruning, neuronal circuit refinement, and tissue homeostasis during early development [2]. These processes are tightly regulated across embryonic and postnatal stages, where microglia exhibit dynamic transcriptional and functional changes in response to the evolving neural environment [1, 3].

During development, microglia transition through multiple functional states, ranging from highly phagocytic and proliferative phenotypes in early stages to more quiescent, homeostatic states in the adult brain [4, 5]. Homeostatic microglia are characterized by expression of canonical markers such as Tmem119 and P2ry12, whereas developmental microglia exhibit elevated expression of genes associated with phagocytosis, lipid metabolism, and stress responses [6, 7]. In addition to these states, emerging evidence highlights the role of oxidative stress pathways and chemokine signaling in shaping microglial behavior, suggesting that microglial identity is regulated by a combination of intrinsic transcriptional programmes and environmental cues [4, 5].

Among these regulatory pathways, the heme oxygenase-1 (*Hmox1*) axis represents a key mediator of cellular responses to oxidative stress [8–11]. *Hmox1*, a downstream target of the nuclear factor erythroid 2-related factor 2 (*NRF2*) signaling, catalyzes the degradation of heme into biologically active metabolites and plays a protective role in maintaining cellular redox balance [4, 5, 8, 9, 11]. In microglia, activation of the *NRF2/Hmox1* pathway has been associated with anti-inflammatory responses, metabolic adaptation, and modulation of cellular stress pathways [12, 13]. However, the precise role of *Hmox1* during normal microglial development remains incompletely understood [13, 14].

Dysregulation of *Hmox1* and related oxidative stress pathways has been implicated in a range of neurological and developmental disorders, including neuroinflammation, neurodegeneration, and perinatal brain injury [4, 13, 15]. Altered microglial activation states during critical development windows may contribute to long-term neurological outcomes, highlighting the importance of understanding how stress-response pathways intersect with microglial maturation [3, 14].

In this study, we investigate the transcriptional landscape of microglial development using pseudobulk RNA-sequencing data spanning embryonic to adult stages. By integrating dimensionality reduction, module scoring, differential expression analysis, and trajectory modelling, we aim to characterize the dynamic transitions between microglial states and to define the role of *NRF2/Hmox1*-associated pathways in shaping these developmental trajectories.

## 3. Materials and Methods

### 3.1 Experimental Design and RNA-Sequencing Data

Publicly available bulk RNA-sequencing data were obtained from Mouse Microglia Bennett et al., 2016, National Center for Biotechnology Information (NCBI) BioProject (accession no. PRJNA30727, doi: 10.1073/pnas.1525528113) [6]. The dataset comprises pseudobulk RNA-sequencing profiles of microglia and myeloid cell populations isolated from the developing mouse cerebral cortex by fluorescence-activated cell sorting (FACS). Twenty-five individual biological replicate samples span five developmental timepoints: embryonic day 17 (E17), postnatal day 7 (P7), postnatal day 14 (P14), postnatal day 21 (P21), and postnatal day 60 (P60). Within each timepoint, cells were sorted by surface expression of Tmem119 into two fractions: a Tmem119^+^ fraction, enriched for mature, homeostatic microglia, and a Tmem119^-^ fraction, enriched for developmentally immature or activation-shifted populations. Importantly, E17 replicates are exclusively Tmem119^-^, as Tmem119^+^ cells were not present at this developmental stage [6]. An additional unsorted microglia/macrophage fraction was collected but excluded from primary analyses, as their heterogeneous transcriptional profiles exerted a strong outlier leverage on principal component axes and obscured biologically meaningful variation within the sorted microglial compartment. Raw expression values for all detected transcripts were imported, and gene symbols were standardised by removing species annotation suffixes from Ensembl gene identifiers. Where multiple probes mapped to the same gene symbol, expression values were collapsed by computing the arithmetic mean across probes. All expression values were log₂-transformed with a pseudocount of 1 [log₂(counts + 1)] to stabilise variance across the dynamic range of the data. Raw counts were assumed to be library-size normalized as part of the pseudobulk processing of the original dataset, and no additional normalization was applied prior to log transformation. Gene Ontogeny (GO) enrichment analysis was not performed, as the analysis focused on hypothesis-driven gene modules rather than unbiased functional enrichment approaches.

### 3.2 Sample Quality Control and Filtering

Sample identifiers were parsed from column headers using a double-underscore delimiter (format: stage sort celltype_rep) to extract fields for developmental stage, sort fraction, cell type, and replicate number. Two categories of samples were excluded before dimensionality reduction and all downstream analyses. First, microglia samples from adults who received intraperitoneal LPS at P60were removed, as this acute inflammatory perturbation falls outside the scope of the developmental trajectory reported here. Second, unsorted microglia/macrophage bulk fractions were excluded because their markedly divergent transcriptional profiles, reflecting a heterogeneous mixture of brain-resident and infiltrating myeloid cells, exerted strong outlier leverage on principal component axes and obscured biologically meaningful variation within the sorted populations. The final analytical dataset comprised 20 samples from five developmental stages across Tmem119^+^ and Tmem119^-^ sort fractions: E17 (n = 3, Tmem119^-^), P7 (n = 4; 2 Tmem119^+^ and 2 Tmem119^-^), P14 (n = 3, Tmem119^+^), P21 (n = 5; 3 Tmem119^+^ and 2 Tmem119^-^), and P60 (n = 5; 3 Tmem119^+^ and 2 Tmem119^-^).

### 3.3 Principal Component Analysis (PCA)

PCA was performed on the filtered and log₂-normalized expression matrix using the prcomp() function in R with scaling parameter enabled (scale. = TRUE). Prior to PCA, genes were pre-filtered to retain only those with variance above the 50th percentile (n=11617) across the 20 clean samples, to reduce noise and focus on high-variance genes. For this analysis, ‘clean samples’ refers to the 20 quality-filtered naïve samples remaining after exclusion of microglia from adults who received intraperitoneal LPS at P60 and unsorted microglia/macrophage fractions. The percentage of variance explained by each principal component was calculated as the squared singular value divided by the total sum of squared singular values. PCA scores for the first two principal components (PC1 and PC2) were retained for all subsequent visualizations and module-scoring analyses. PCA scatter plots were generated to characterize sample relationships by developmental stage; sort population (Tmem119^+^ versus Tmem119^-^); dominant microglial transcriptional programme; and individual gene expression (feature plots). For feature plots, expression values were color-clamped to the 5th-95th percentile range to improve contrast without distorting extreme values. Each developmental stage was assigned a fixed color (E17 = red, P7 = orange, P14 = yellow, P21 = green, P60 = blue), and sort fractions were distinguished by point shape (filled circle for Tmem119^+^; triangle for Tmem119^-^).

### 3.4 Microglial Programme Module Scoring

Four transcriptional programme modules were defined a priori based on established microglial biology and curated gene signatures [6, 7, 13]. (i) The homeostatic programme comprised *Tmem119, P2ry12, Siglech,* and *Tgfbr1*, canonical markers of surveilling, ramified microglia in the healthy adult brain [6, 7, 16]. (ii) The phagocytic/disease-associated programme comprised *Gpnmb, Igf1, Csf1, Lgals3, Fabp5, Spp1,* and *Apoe*, genes enriched in active phagocytic and disease-associated microglial states [17–20]. (iii) *NRF2/Hmox1* oxidative stress response programme comprised *Hmox1, Nqo1, Gclc, Gclm, Srxn1, Txnrd1, Slc7a11, Fth1*, and *Ftl1*, effectors of the canonical *NRF2* antioxidant response [4, 5, 13, 21, 22]. (iv) The *Apoc1*-associated programme comprised *Apoc1* and *Lyz2*, genes associated with lipid metabolism and myeloid cell identity [14]. Module scores were computed on a z-scored expression matrix to place all four programmes on a comparable scale. Briefly, each gene’s expression vector was standardized across all 20 samples to a mean of zero and a standard deviation of one, such that genes with constitutively high abundance (e.g. *Tmem119*, log_2_ expression ≈ 10) did not numerically dominate genes with lower baseline expression (e.g. *Spp1*, log_2_ expression ≈ 3). The module score for each sample was then calculated as the arithmetic mean of the z-scored values across all member genes of a given programme.

### 3.5 Differential Expression Analysis

Differential expression (DE) analysis was performed on log₂-transformed expression values using a gene-wise two-sample Welch’s t-test, implemented via the base R t.test() function, which does not assume equal variance between groups. Welch’s t-test was used due to pseudobulk log_2_ normalized values displaying a Gaussian distribution and relatively small sample size rendering standard count- based models (e.g., DESeq2) less appropriate. Multiple testing correction was applied genome-wide using the Benjamini-Hochberg (BH) false discovery rate (FDR) procedure via p.adjust(method = "BH"). Genes with |log2FC| > 1 and BH raw p < 0.05 were considered differentially expressed [14, 23]. Two independent DE comparisons were conducted. First, Tmem119^+^ versus Tmem119^-^ sorted microglia were compared across all developmental ages. Second, developmentally early samples (E17 combined with P7) were compared against mature microglia (P60) to characterize the transcriptional changes accompanying microglial maturation. Results were visualized as volcano plots in which the x-axis represents log₂ fold change and the y-axis represents -log₁₀(raw p-value).

### 3.6 Hierarchical Clustering and Heatmap Visualization

Heatmaps were generated using the pheatmap R package. Expression values were row-scaled (z-scored per gene) prior to display. Genes were ordered by hierarchical clustering of rows (complete linkage, Euclidean distance), while columns were ordered by developmental stage and sort population without column clustering, to preserve the developmental axis. A combined marker heatmap was constructed from genes encompassing all four microglial programme gene sets together with additional myeloid marker genes. A separate heatmap was generated for chemokine receptor genes.

### 3.7 Dot Plot Visualization of Marker Gene Expression

Dot plots were used to summarize marker gene expression across developmental stages. For each gene-stage combination, two summary statistics were calculated: the average log₂-transformed expression across all samples at that stage, and the percentage of samples with detectable expression, defined as at least one count [log₂(count + 1) > 0]. These statistics were encoded as dot color (average expression, light grey to magenta gradient) and dot size (percentage of samples expressing the gene).

### 3.8 Developmental Trajectory and Pseudotime Analysis

To model the transcriptional continuum of microglial maturation, a principal curve was fitted to PCA coordinates derived from a trajectory-specific subset of samples comprising three biologically representative developmental stages: embryonic (E17), weaning age (P21), and young adulthood (P60), using the principal_curve() function from the princurve R package (v2.1.6), with a smooth spline smoother (df=4). A principal curve approach was selected over graph-based pseudotime methods (e.g., Monocle, Slingshot) due to the low sample size and pseudobulk structure of the dataset, which limit the applicability of single-cell trajectory inference frameworks [24]. The arc-length parameter (λ) along the fitted curve was linearly rescaled to a pseudotime range of 0-25, chosen to provide sufficient numerical resolution to visualize gradual transcriptional transitions across developmental stages while remaining interpretable. Module scores for the homeostatic, phagocytic, and *NRF2/Hmox1* oxidative stress response programmes were computed on the trajectory-specific z-scored expression matrix and plotted against pseudotime with LOESS smoothing curves overlaid (span = 0.75, 95% confidence intervals shown).

### 3.9 Chemokine Receptor Expression Analysis

Six microglia chemokine receptors (*Cxcr1, Ccr5, Cxcr4, Ccr3, Ccr4, Cxcr2*) were selected a priori based on their previously identified or putative roles in surveillance, recruitment and inflammatory signaling to determine their expression across development [25–30]. Of these, *Cxcr1* showed zero detectable expression across all 20 samples and was therefore excluded from heatmap clustering and module scoring, though it is retained in boxplot and feature plot panels for transparency. The remaining five receptors were characterized across developmental stages and sort fractions using five complementary visualization approaches: per-gene box plots, a dot plot, individual PCA feature plots, a z-scored module score overlay on the PCA plot, and a row-scaled heatmap.

### 3.10 Targeted Gene Expression Visualization

Expression of biologically prioritized gene panels was visualized using per-gene box plots stratified by developmental stage and sort fraction, with individual data points overlaid. Three panels were examined: (i) transcriptional expression of common flow cytometry panel markers (P2ry12, Xcr1, Cx3cr1, Cxcr3, Cd163, Cd11bc, Cd45), (ii) *NRF2/Hmox1* oxidative stress programme genes, and (iii) cluster-defining marker genes (*Slco2b1, Ccl12, Apoe, Ccdc152, Cdkn1a, Spp1, Fabp5, Hmox1, Arg1, Apoc1*, and *Ifi27l2a*). *Hmox1* appears in both panels (ii) and (iii), reflecting its dual identity as a canonical *NRF2* target and a cluster-defining marker [14].

### 3.11 Software and Statistical Computing

All analyses were performed in R (version 4.4.2). Data manipulation and visualization used the tidyverse suite (version 2.0.0). Principal curve fitting used the princurve package (version 2.1.6). All stochastic steps used a random seed of 42 to ensure reproducibility. For differential expression analyses, statistical significance was defined as raw p-value < 0.05 combined with |log_2_FC| > 1, as described in Section 3.5. BH-adjusted p-values (FDR) were computed genome-wide and are reported in the supplementary tables, but volcano plot thresholds are applied to raw p-values given the exploratory nature of the visualization. Analysis code is available within the supplementary data document.

## 4. Results

### 4.1 Transcriptional profiling of naïve mouse microglia across brain development

To characterize the transcriptional identity of mouse microglia across brain development, we analyzed bulk RNA-sequencing data (accession no. PRJNA30727, doi: 10.1073/pnas.1525528113) [6] from sorted microglial populations spanning five developmental stages: embryonic day 17 (E17), postnatal days 7 (P7), 14 (P14), 21 (P21), and 60 (P60). Samples were sorted by Tmem119 expression into Tmem119^+^ and Tmem119^-^ populations to distinguish mature homeostatic microglia from other myeloid cells [6, 31]. Unsorted microglia-macrophage samples and microglia samples from adults who received intraperitoneal LPS at P60 were excluded from quantitative analyses to avoid contamination by non-microglial myeloid cells and to focus on naïve states in the absence of inflammation, respectively. Following quality filtering, the final dataset comprised 20 samples, including five developmental stages and two sort fractions.

Principal component analysis (PCA) of the top 50% most variable genes separated samples according to developmental stage along PC1 and PC2, demonstrating a clear transcriptional trajectory from early embryonic to mature adult microglia **(figure 1A).** Tmem119^+^ and Tmem119^-^ populations were further resolved by point shape. Feature plots on the PCA embedding confirmed that the microglia-enriched marker *Sparc* **(figure 1B)** was preferentially expressed in Tmem119^+^ populations, while the border-associated macrophage marker *Mrc1* **(figure 1C)** was enriched in Tmem119^-^ fractions The homeostatic marker P2ry12 displayed strong stage and sort-dependent expression, with highest levels in mature Tmem119^+^ microglia **(figure 1D).**

**Figure 1:**
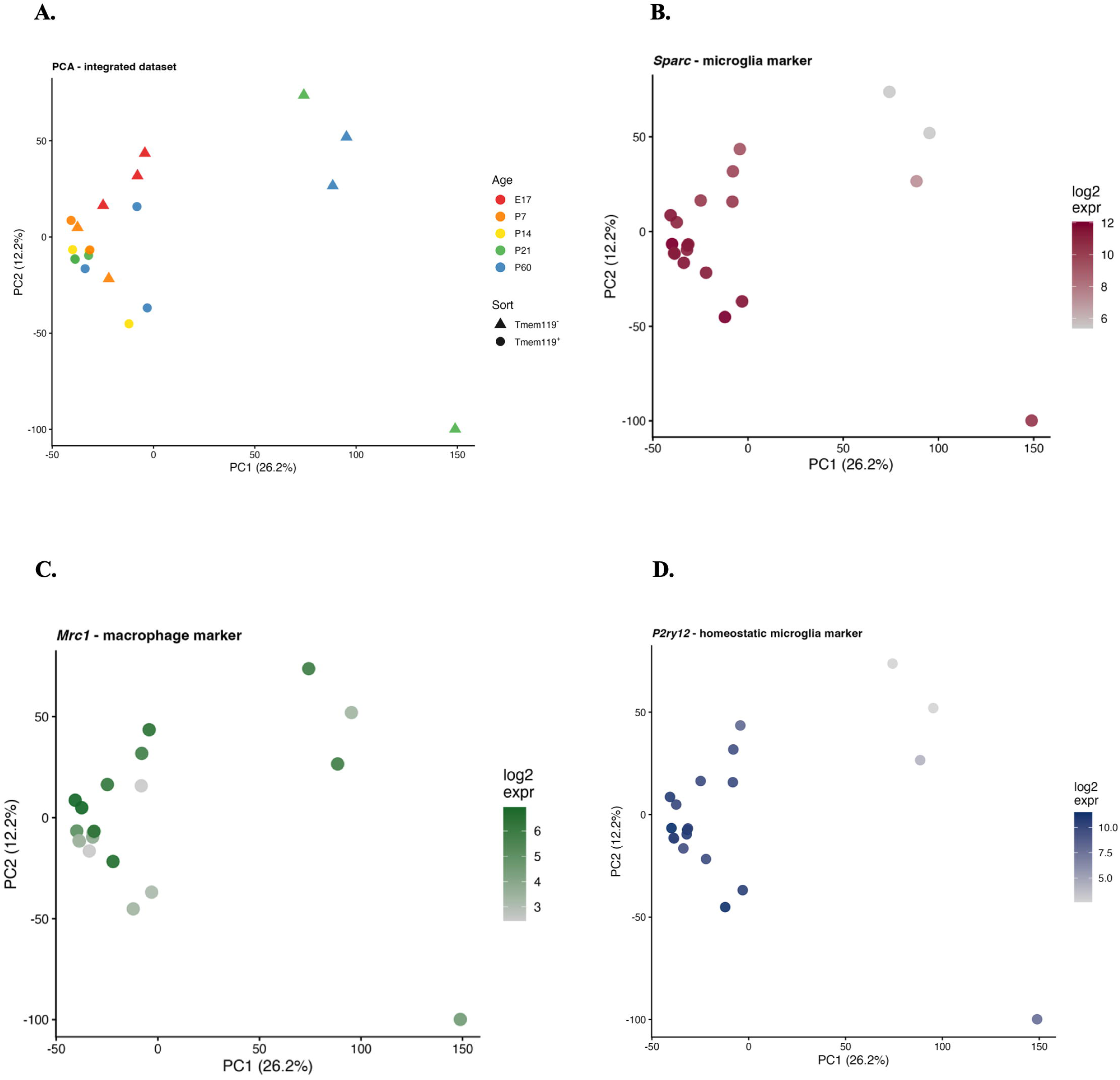
Developmental Transcriptional Landscape of Mouse Microglia. **Figure 1A.** PCA by age showing developmental stage separation along principal components. Each point represents one sample; colors indicate developmental stage (E17=red, P7=orange, P14=yellow, P21=green, P60=blue); shapes distinguish Tmem119^+^ (circle) from Tmem119^-^ (triangle) fractions. **Figure 1B**. PCA feature plots showing expression of *Sparc* (microglia-enriched marker). **Figure 1C**. *Mrc1* (border-associated macrophage marker), confirming sort-fraction separation. **Figure 1D**. PCA feature plot of *P2ry12* homeostatic marker expression, showing highest levels in mature Tmem119^+^ microglia.

### 4.2 Developmental regulation of microglial programme marker genes

To characterize the temporal dynamics of established microglial marker genes, we examined expression across development using dot plots encoding average expression and the proportion of samples with detectable transcripts [6, 14]. Broad myeloid class markers spanning the homeostatic, phagocytic, *NRF2/Hmox1*, and *Apoc1*-like programmes, together with *Ccr2*, *Mrc1*, and *Lyve1*, revealed distinct developmental expression patterns **(figure 2A).** Homeostatic markers (*Tmem119, P2ry12, Siglech, Tgfbr1*) increased progressively with age, consistent with the acquisition of a mature microglial identity. In contrast, phagocytic and *NRF2*-associated genes showed comparatively higher expression at early developmental stages.

**Figure 2:**
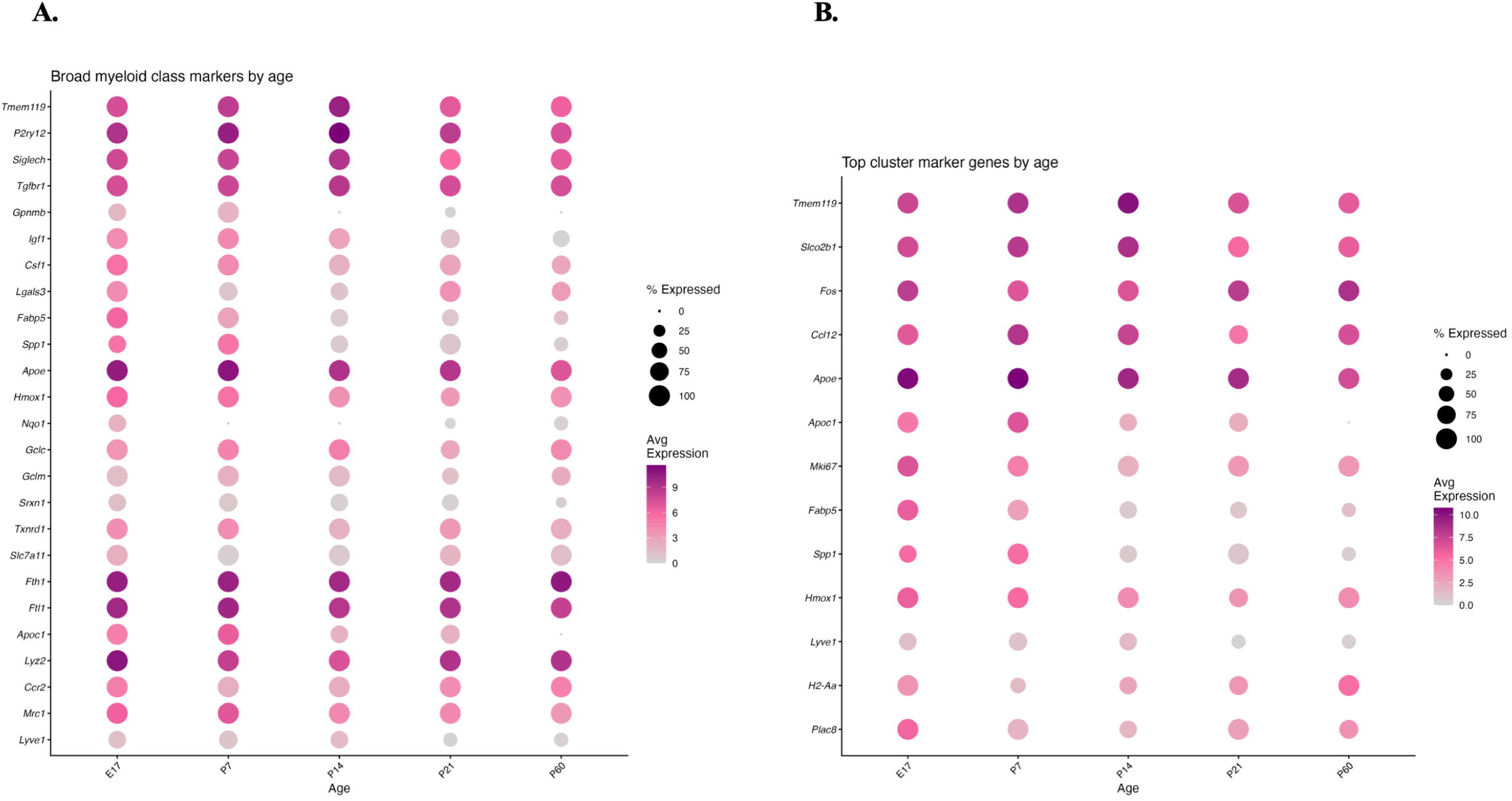
Developmental Marker Dynamics. **Figure 2A**. Dotplot showing myeloid class markers by age. **Figure2B.** Dotplot showing the top cluster marker genes by age.

Analysis of top cluster marker genes including *Slco2b1, Fos, Ccl12, Apoe, Apoc1, Mki67, Fabp5, Spp1, Hmox1, Lyve1, H2-Aa,* and *Plac8* further delineated distinct cellular states across development **(figure 2B).** Proliferation marker *Mki67* was detected predominantly at early timepoints, while lipid-associated genes including *Fabp5* and *Apoe* showed enrichment in early postnatal microglia. **(figure 2C).**

### 4.3 Z-scored module scoring identifies four distinct microglial transcriptional programmes in the brain

To systematically quantify the relative activity of distinct microglial states, we computed z-scored module scores for four transcriptional programmes: Homeostatic, Phagocytic, *NRF2/Hmox1*, and *Apoc1*-like [14]. Z-scoring was applied to each gene across samples prior to module averaging to ensure that programmes with high-abundance genes (e.g. *Tmem119*, log2 expression ≈ 10) did not numerically dominate those with lower-abundance genes (e.g. *Spp1*, log2 expression ≈ 3).

PCA of the full dataset, colored by dominant programme, demonstrated clear spatial separation between Homeostatic and Phagocytic/*NRF2* programme states **(figure 3A).** Violin plots of z-scored module scores across developmental stages showed that the Homeostatic programme was progressively dominant at later timepoints (P21, P60), while the Phagocytic and *NRF2/Hmox1* programmes showed higher relative scores at early stages (E17, P7) **(figure 3B).** The *Apoc1*-like programme was detected at low levels throughout development. Programme frequency analysis confirmed these developmental trends: the proportion of samples classified as Homeostatic increased from E17 to P60, while the proportion of Phagocytic-dominant samples declined with age **(figure 3C and 3D).** Faceted PCA plots across each developmental stage illustrated progressive spatial compaction of the sample cloud as microglia converged on a mature homeostatic state by P60 **(figure 3E).**

**Figure 3:**
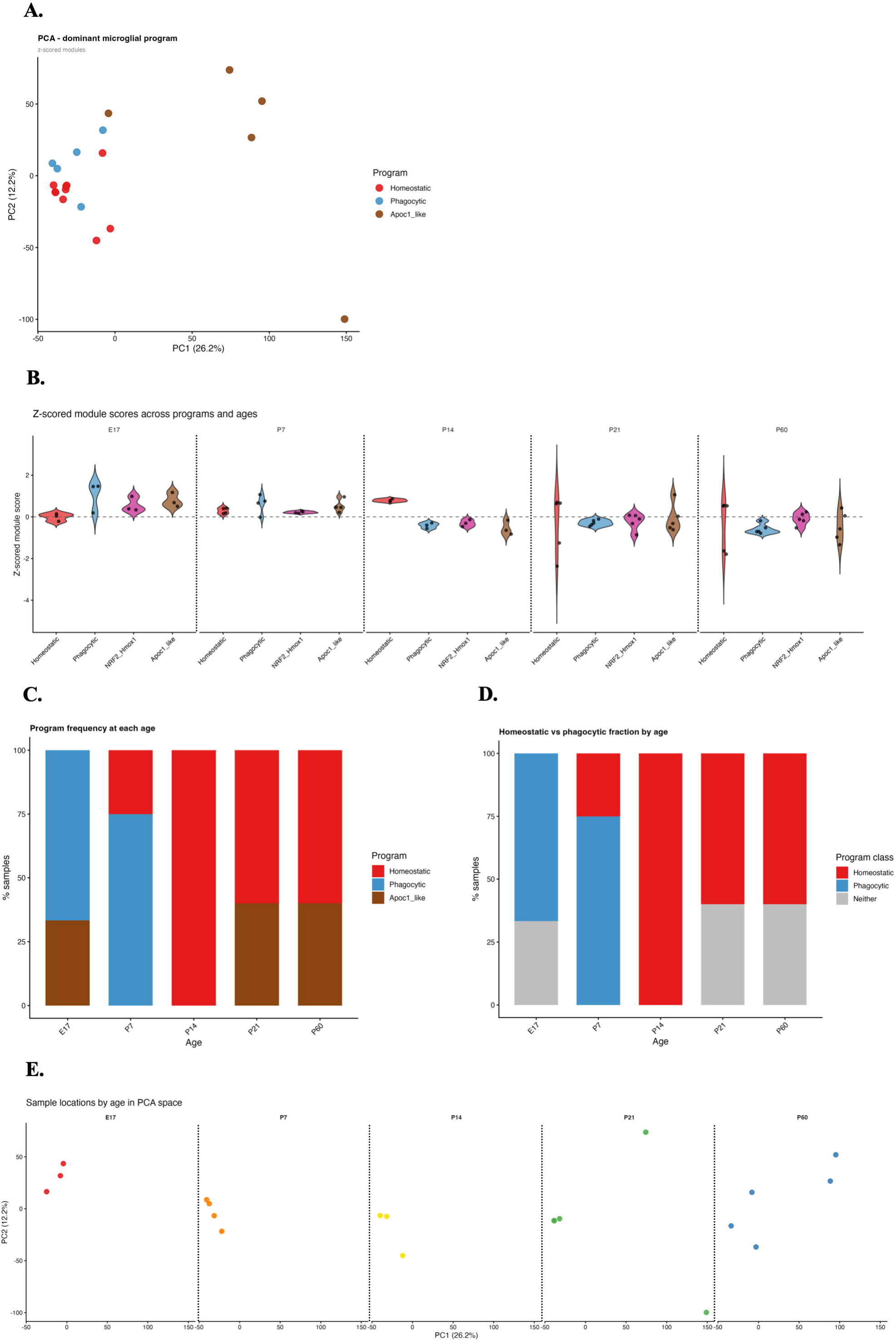
Microglial Programme Architecture Across Development. **Figure 3A**. PCA colored by dominant microglial transcriptional programme, showing spatial separation of Homeostatic from Phagocytic/NRF2 states. **Figure 3B**. Violin plots of z-scored module scores across developmental stages for all four transcriptional programmes. **Figure 3C**. Programme frequency by age, stacked bar chart showing percentage of samples assigned to each dominant transcriptional programme at each developmental stage. **Figure 3D**. Homeostatic vs. phagocytic fraction, two-class classification showing the developmental shift from phagocytic to homeostatic dominance. **Figure 3E**. PCA faceted by age, illustrating progressive spatial compaction as microglia mature toward a homeostatic phenotype by P60.

### 4.4 Module score gradients across principal component space

Module score gradients overlaid on the PCA space confirmed high Homeostatic scores in mature Tmem119^+^ P60 microglia **(figure 4A)**, elevated Phagocytic scores in early-stage samples **(figure 4B)**, and *NRF2/Hmox1* activity distributed across intermediate developmental stages **(figure 4C)**, consistent with a transitional regulatory role for oxidative stress signaling during microglial maturation [4, 13].

**Figure 4:**
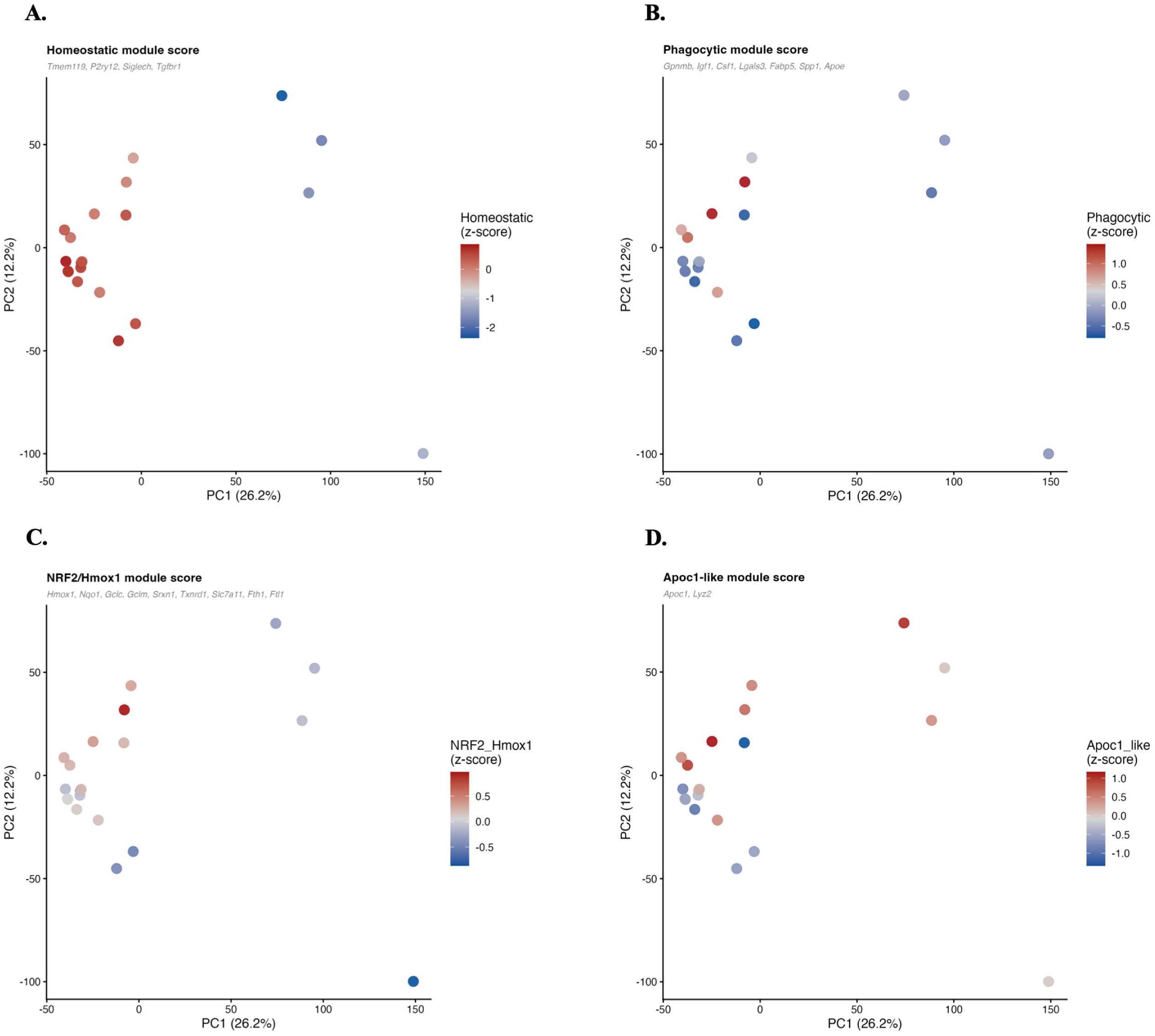
Functional Programme Gradients in PCA Space. **Figure 4A**. Homeostatic module score overlaid on PCA space, showing enrichment in mature P60 Tmem119^+^ microglia. **Figure 4B**. Phagocytic module score overlaid on PCA space, showing enrichment at early developmental timepoints. **Figure 4C**. *NRF2/Hmox1* module score overlaid on PCA space, with activity concentrated at intermediate developmental stages. **Figure 4D**. *Apoc1* overlaid on PCA space, showing enrichment at early developmental timepoints.

### 4.5 Differential expression analysis reveals transcriptional signatures of microglial identity and maturation

To identify gene expression differences between microglial populations, we performed two independent differential expression analyses using gene-wise two-sample t-tests with Benjamini-Hochberg FDR correction. Genes meeting both an absolute log2 fold-change threshold greater than 1 and a nominal p-value less than 0.05 were considered differentially expressed. Comparing Tmem119^+^ versus Tmem119^-^ sorted populations, numerous genes were significantly upregulated in Tmem119^+^ microglia, including established homeostatic markers **(figure 5A).** Prioritized genes from the homeostatic programme (*Tmem119, P2ry12, Siglech*) and the common flow cytometry panel markers (*P2ry12, Cx3cr1*) were consistently enriched in the Tmem119^+^ fraction. Genes associated with border-associated macrophages and monocyte-derived cells were among those upregulated in the Tmem119^-^population. The comparison of early developmental (E17 and P7 combined) versus mature (P60) microglia identified broad transcriptional remodeling across maturation **(figure 5B).** Phagocytic programme genes including *Spp1, Fabp5, Apoe,* and *Gpnmb* were significantly upregulated in early developmental microglia, consistent with their heightened phagocytic and lipid-processing activity during brain development. Homeostatic genes (*Tmem119, P2ry12, Siglech*) and *NRF2* target genes (*Hmox1, Nqo1*) showed differential expression between developmental and mature states **(figure 5B).**

**Figure 5:**
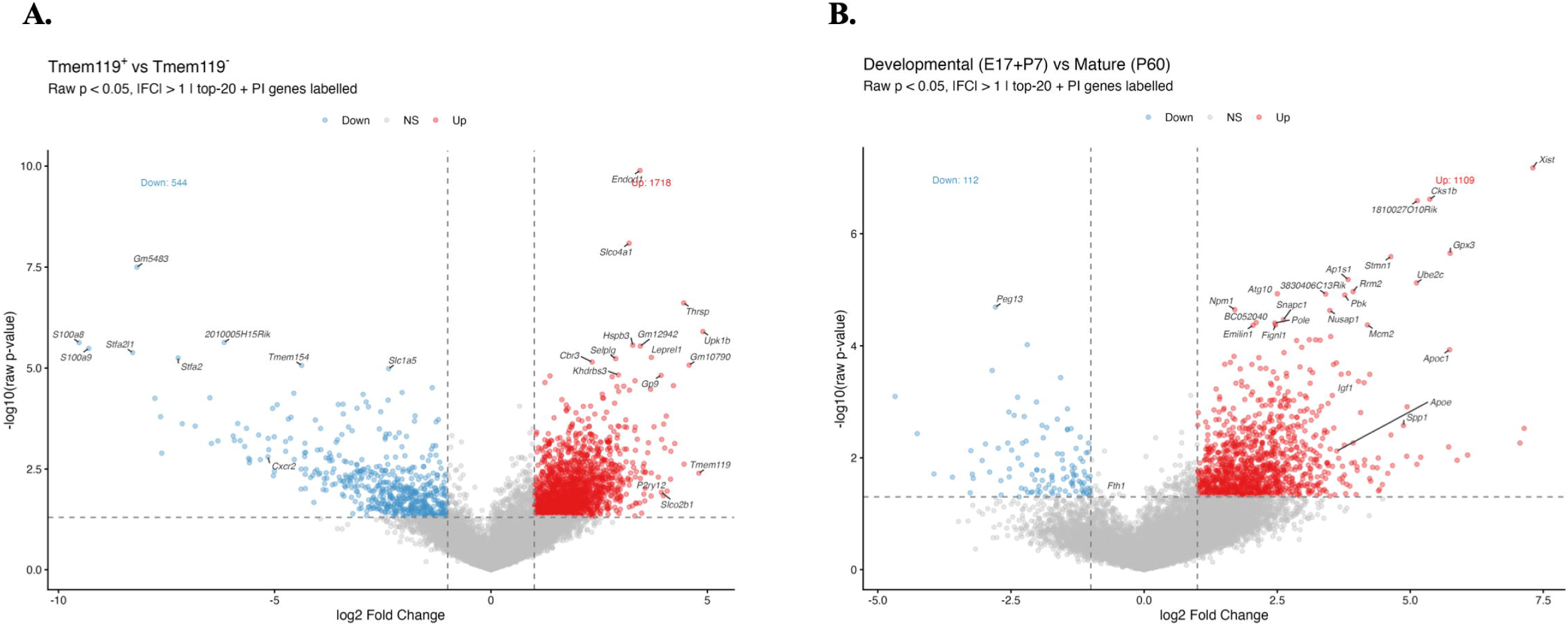
Differential Expression Landscape. Significantly upregulated genes (|log₂FC| > 1, p < 0.05) are shown in red while downregulated genes are displayed in blue; top 20 most significant and biologically prioritized genes are labeled. **Figure 5A**. Volcano plot of Tmem119^+^ vs. Tmem119^-^differential expression. **Figure 5B**. Volcano plot of early developmental (E17 + P7) vs. mature (P60) microglia differential expression. Phagocytic genes are upregulated in early microglia; homeostatic genes predominate at P60.

### 4.6 Heatmap analysis of key microglial marker genes

Unsupervised hierarchical clustering of up to 30 key microglial marker genes across the 20 developmental samples revealed co-regulated gene clusters corresponding to known microglial states [14, 23]. Samples were ordered by developmental stage and sort population without column clustering. Homeostatic marker genes (*Tmem119, P2ry12, Siglech, Tgfbr1*) formed a co-expression cluster with high expression in P60 Tmem119^+^ microglia. Phagocytic genes (*Gpnmb, Spp1, Fabp5, Apoe*) and *NRF2* pathway genes (*Hmox1, Nqo1, Gclc*) formed distinct clusters with elevated expression at earlier stages (E17-P7) and in Tmem119^-^ populations **(figure 6A).** Chemokine markers *Cxcr2* and *Cxcr4* were predominant in Tmem119^-^ myeloid cells at P60, while *Ccr4* was present in Tmem119^-^ microglia at E17 **(figure 6B).**

**Figure 6:**
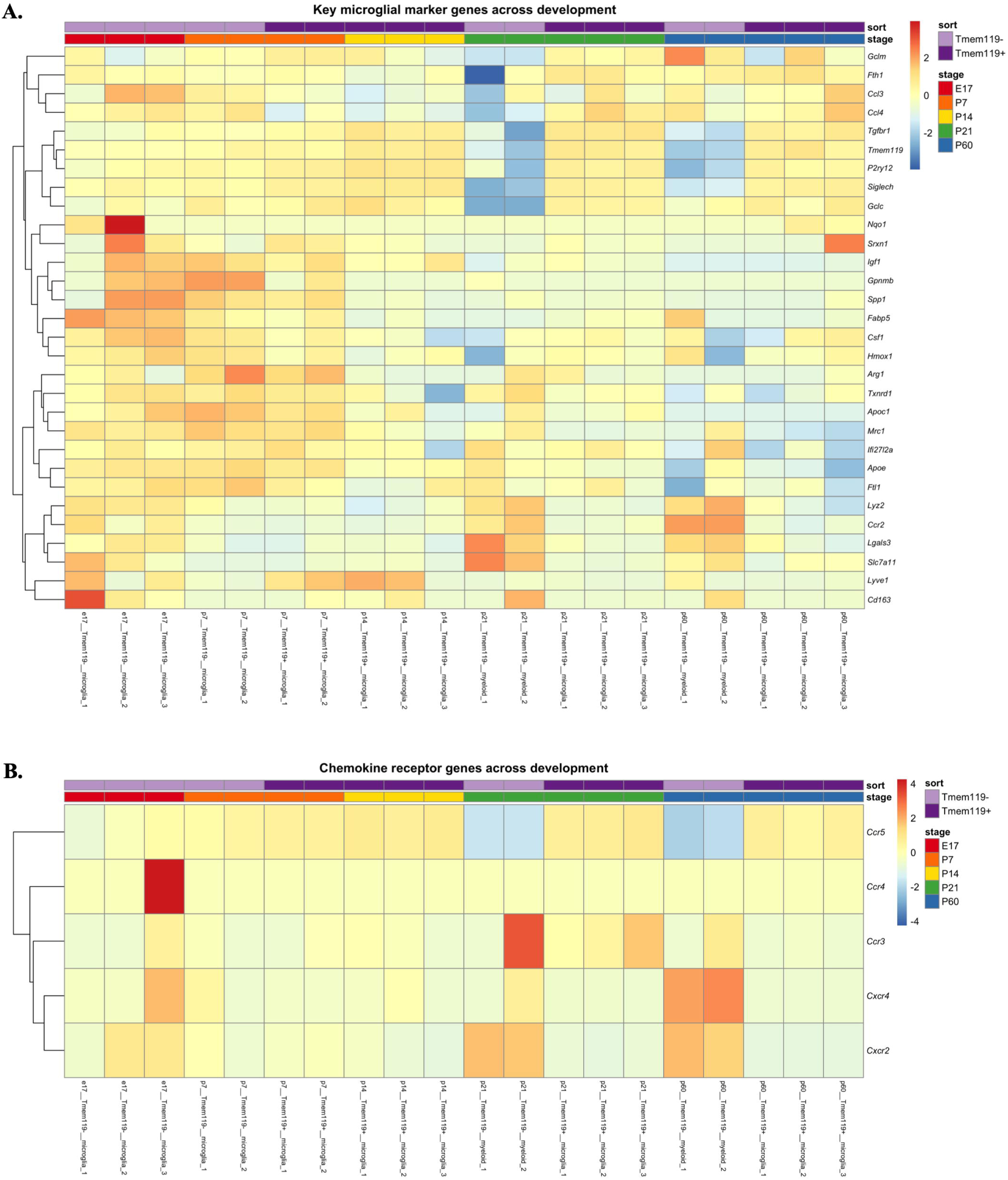
Heatmap of Microglial State Genes. **Figure 6A**. Heatmap of key microglial marker genes across the 20 developmental samples, ordered by developmental stage and sort fraction. Row z-scores shown. **Figure 6B**. Heatmap of chemokine receptor gene expression across developmental stages, with zero-variance genes excluded.

### 4.7 Pseudotime trajectory analysis models a continuous developmental axis from E17 to P60

To model microglial maturation as a continuous transcriptional process, we fitted a principal curve to PCA coordinates derived from E17, P21, and P60 Tmem119^+^ microglia (with all E17 samples included regardless of sort, as E17 microglia are exclusively Tmem119^-^). The resulting pseudotime axis (scaled 0-25) was anchored such that E17 samples occupied low pseudotime values and P60 samples occupied high pseudotime values **(figure 7A-B).** Plotting z-scored module scores along the pseudotime axis revealed reciprocal dynamics between microglial programmes across maturation. The Phagocytic module score declined progressively from E17 (low pseudotime) to P60 (high pseudotime), consistent with the well-established transition from a phagocytic developmental microglia state to a quiescent homeostatic state in the naive adult brain. Conversely, the Homeostatic module score increased along the pseudotime axis. The *NRF2/Hmox1* module showed intermediate dynamics, with elevated scores at mid-developmental stages before declining in fully mature microglia. LOESS smoothing curves confirmed these trajectories with 95% confidence intervals **(figure 7C).**

**Figure 7:**
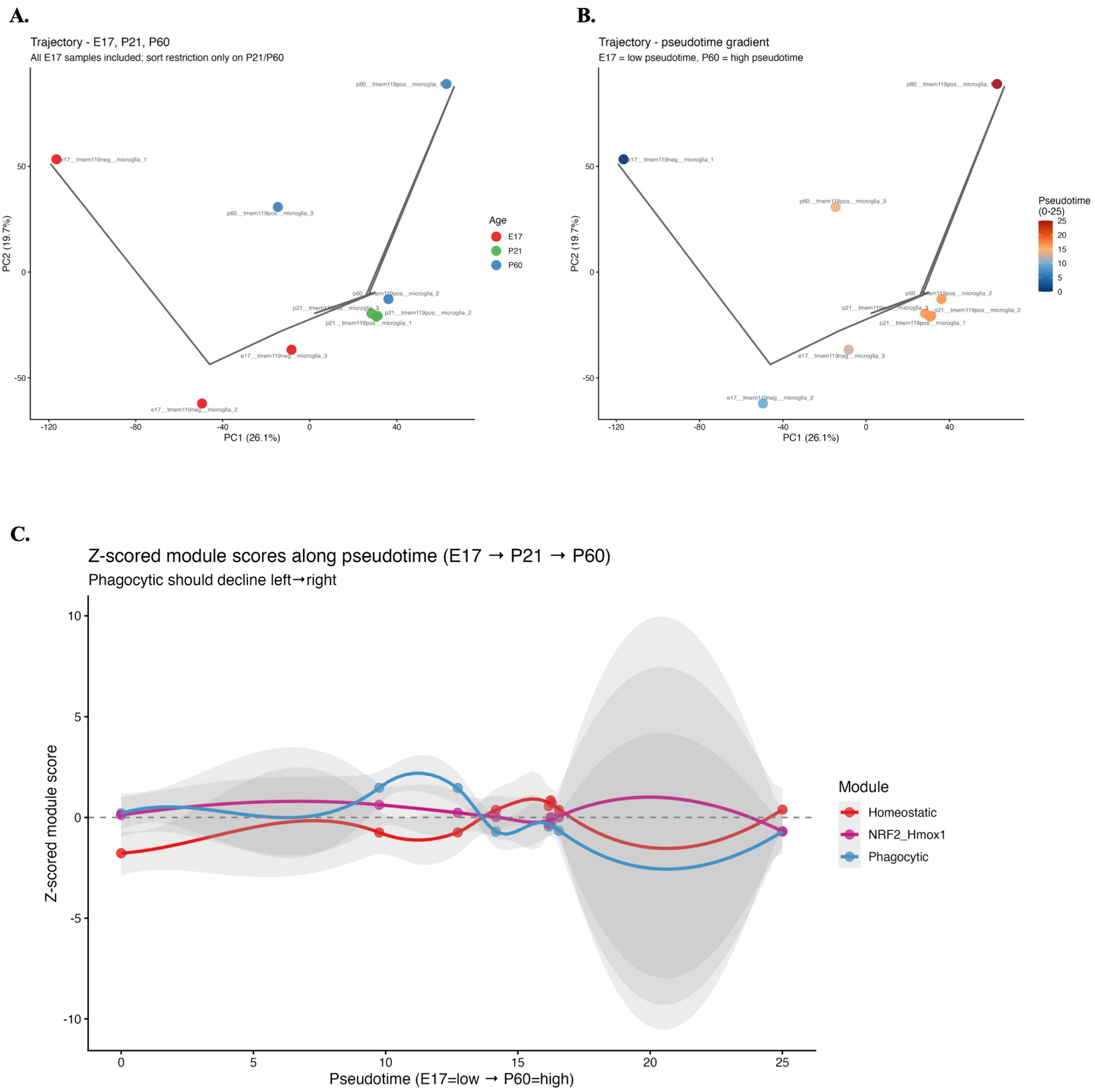
Developmental Trajectory and Pseudotime Analysis. **Figure 7A**. Principal curve fitted to E17, P21, and P60 samples in PCA space, colored by developmental age. The curve captures the continuous developmental trajectory. **Figure 7B**. Same principal curve colored by pseudotime value (scaled 0-25), anchored with E17 at low pseudotime and P60 at high pseudotime. **Figure 7C**. Module scores plotted along the pseudotime axis with LOESS smoothing. The Phagocytic programme declines, the Homeostatic programme rises, and NRF2/Hmox1 peaks at intermediate stages.

### 4.8 Chemokine receptor expression across microglial development

The expression of six chemokine receptors (*Cxcr1, Ccr5, Cxcr4, Ccr3, Ccr4, Cxcr2*) was examined across developmental stages and sort populations. Among these, *Cxcr1* was not detected in the dataset (zero expression across all 20 samples) and was therefore excluded from heatmap clustering. *Ccr5* and *Cxcr2* showed the highest average expression levels across developmental stages **(figure 8A).** *Cxcr4*, a receptor important for microglial migration and brain development, showed detectable expression particularly at early timepoints (E17 and P7) **(figure 8A).** The z-scored chemokine receptor module score was not strongly associated with either the Tmem119^+^ or Tmem119^-^ sorted populations, suggesting these receptors are expressed broadly across myeloid cell subsets **(figure 8 B and C)**. Microglia chemokine receptors across development and in PCA space are shown in **supplementary figure 1A and 1B**, respectively.

**Figure 8:**
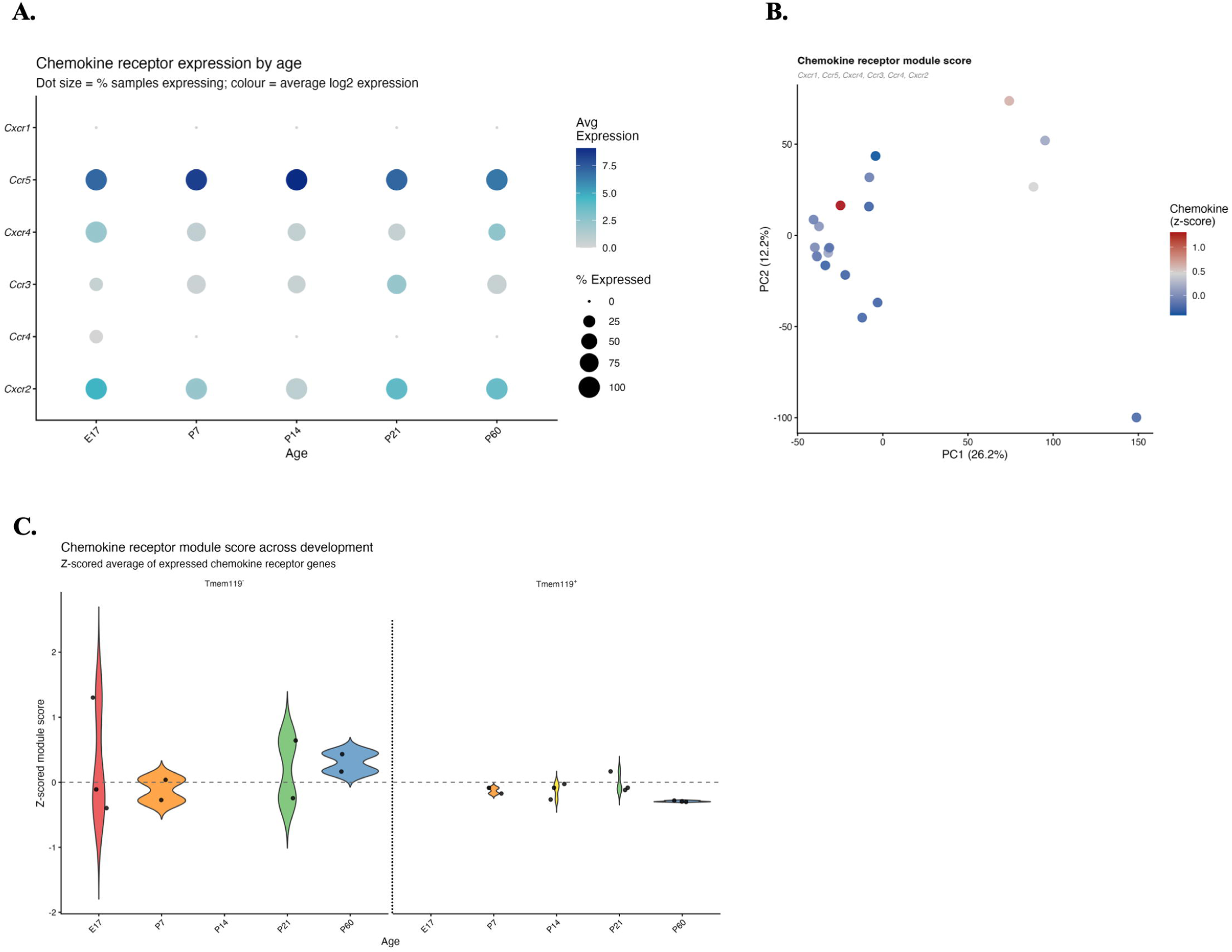
Chemokine Receptor Programmes. **Figure 8A**. Dot plot of chemokine receptor gene expression across developmental stages, showing average expression (color) and percentage of expressing samples (size). **Figure 8B**. Chemokine receptor module score overlaid on PCA space, showing broad distribution across myeloid populations. **Figure 8C**. Violin plots of chemokine receptor expression across developmental stages and Tmem119^-^ and Tmem199^+^ sort fractions.

### 4.9 Expression of biologically prioritised gene panels across development

Boxplot analysis of three prioritized gene panels revealed distinct developmental regulation. Flow cytometry panel markers (*P2ry12, Xcr1, Cx3cr1, Cxcr3, Cd163, Cd11bc, Cd45*) showed stage-and sort-dependent expression consistent with their utility as sorting and identity markers **(figure 9A)**. *P2ry12* and *Cx3cr1* were most highly expressed in mature Tmem119^+^ microglia, while *Cd163* was enriched in Tmem119^-^ populations. *NRF2/Hmox1* pathway genes showed variable expression across development, with *Hmox1* displaying elevated expression at early developmental stages, consistent with heightened oxidative stress responses in developing brain tissue **(figure 9B)**. Cluster-defining marker gene expression *(Apoc1, Apoe, Arg1, Ccdc152, Ccl12, Cdkn1a, Fabp5, Hmox1, Ifi27l2a, Slco2b1, Spp1)* **(figure 9C)** and a combined dot plot integrating all three panels confirmed that these gene sets capture complementary aspects of the transcriptional landscape of developing microglia **(figure 9D)**.

**Figure 9:**
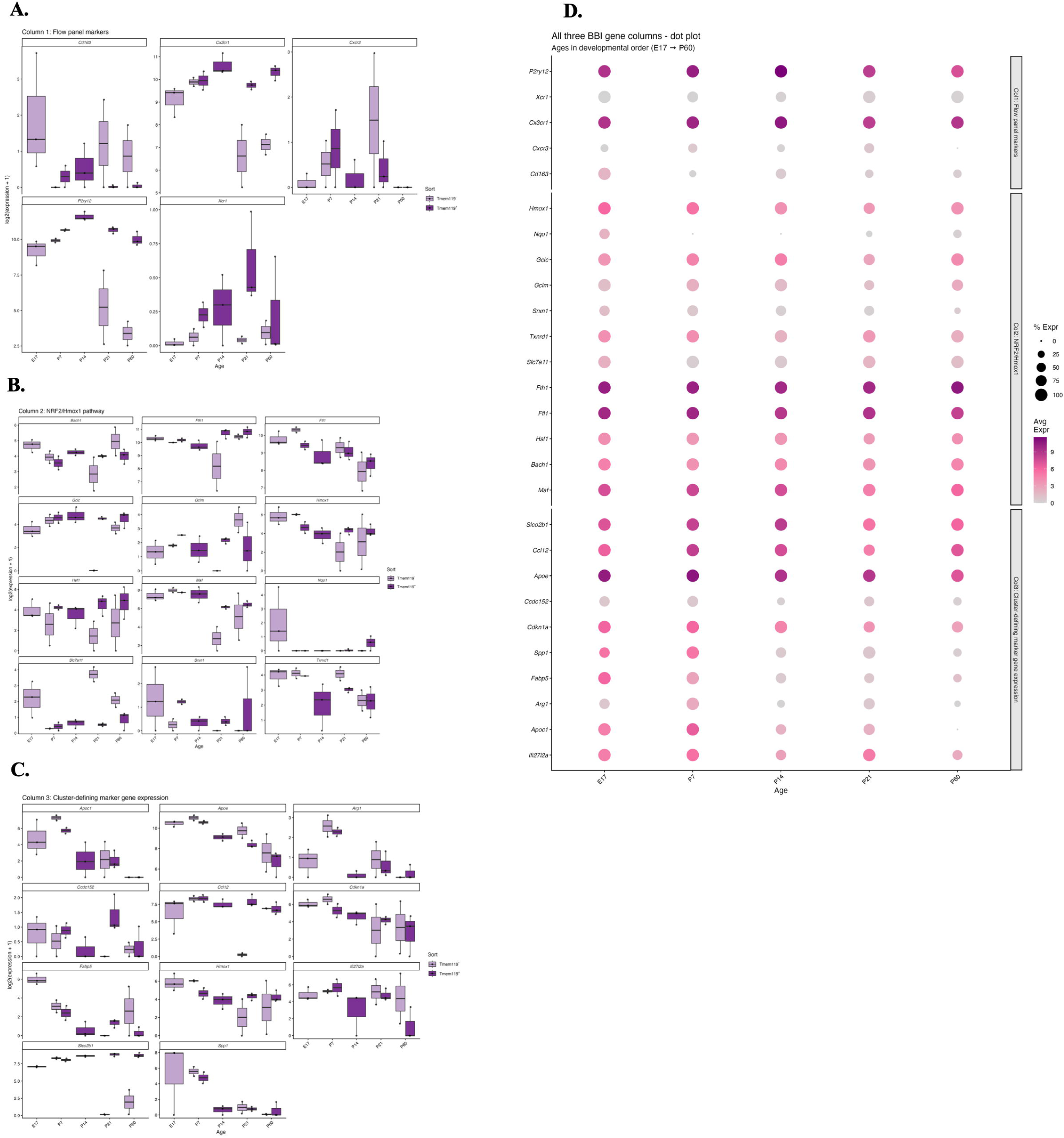
Targeted Gene Panel Validation. **Figure 9A**. Boxplots of flow panel markers *(CD163, Cx3cr1, Cxcr3, P2ry12, Xcr1)* stratified by developmental stage and sort fraction. **Figure 9B**. Boxplots of NRF2/Hmox1 oxidative stress pathway genes *(Bach1, Fth1, Ftl1, Gclc, Gclm, Hmox1, Hsf1, Maf, Nqo1, Slc7a11, Srxn1, Txnrd1)* across development. **Figure 9C**. Boxplots of cluster-defining marker gene expression *(Apoc1, Apoe, Arg1, Ccdc152, Ccl12, Cdkn1a, Fabp5, Hmox1, Ifi27l2a, Slco2b1, Spp1)* stratified by developmental stage and sort fraction. **Figure 9D**. Integrated visualization of all biologically prioritized gene panels across developmental stages.

## 5. Discussion

In this study, we characterized the transcriptional dynamics of microglial development across the embryonic and postnatal stages. Our analyses reveal a clear progression from early phagocytic to a mature homeostatic phenotype, accompanied by coordinated changes in gene expression programmes.

Notably, *NRF2/Hmox1*-associated activity exhibited stage-specific dynamics, suggesting a potential regulatory role during microglial maturation in the brain. Early developmental microglia (E17-P7) were characterized by elevated expression of genes associated with phagocytosis, lipid metabolism, and cellular stress responses, consistent with their established roles in synaptic pruning and tissue remodeling [1, 3]. In contrast, mature microglia (P60) displayed strong enrichment of homeostatic markers, reflecting the acquisition of a stable surveillance phenotype [2]. These suggest that *NRF2*-driven oxidative stress responses and phagocytic activation represent functionally separable but co-occurring microglial states, particularly under conditions of elevated metabolic demand such as those present in the developing brain [2].

Consistent with these findings, PCA visualization demonstrated a clear age-dependent organization of samples, with early-stage microglia occupying a broader transcriptional space and mature microglia converging into a more compact cluster. This pattern is consistent with previous reports that developmental microglia exhibit greater transcriptional heterogeneity compared to their adult counterparts [1, 3]. Module-based analysis identified distinct transcriptional programmes corresponding to homeostatic, phagocytic, oxidative stress-responsive, and lipid-associated microglial states. This four-programme architecture aligns with the multi-state model of microglial identity observed during development [14]. The phagocytic programme was predominantly active at early stages (E17 and P7), supporting its established role in synaptic pruning and apoptotic cell clearance during postnatal brain development [1, 3], while the homeostatic programme became dominant with maturation, consistent with the well-characterised acquisition of a surveilling, ramified phenotype in the adult brain [1].

The *NRF2/Hmox1*-associated programme showed intermediate activation, suggesting a potential transitional role between developmental and mature states in the naïve cerebral cortex. The observed dynamics suggest that oxidative stress signaling may play a critical role in regulating microglial state transitions. Elevated *Hmox1* expression at early stages may reflect increased metabolic and oxidative demands during active tissue remodeling, while its decline in mature microglia is consistent with reduced cellular stress in the healthy adult brain [2, 4, 5, 14]. Our findings defining the *NRF2/Hmox1*- associated programme at naïve state is important because upregulation of *NRF2/Hmox1* programme genes (*Nqo1, Slc7a11, Ftl1*) are associated with inflammation, lipid peroxidation and ferroptosis in microglia [13]. Recent advances in single-cell transcriptomics have expanded our understanding of microglial heterogeneity, revealing multiple functionally distinct subtypes across development and disease [2, 3, 32]. Our findings align with these studies, supporting a model in which microglial identity is not static but evolves along a continuum of transcriptional states influenced by developmental and environmental factors [1–3, 32].

Chemokine receptors represent an additional layer of regulatory complexity in microglial development [33–36]. Among the receptors examined, *Cxcr4* showed detectable expression particularly at early developmental timepoints, consistent with its established role in microglial migration, proliferation, and brain colonization during embryonic development [30, 35, 36]. *Cxcr4* signaling via its ligand CXCL12/SDF-1 has been shown to regulate microglial positioning and survival in the developing brain, suggesting that chemokine-mediated cues contribute to the spatial organization of the early microglial niche [36]. Literature suggests that *Ccr5* acts as the primary receptor for anti-inflammatory differentiation and phagocytic function in microglia under inflammatory conditions [37] whereas *Cxcr2* are expressed by activated and inflammatory microglia [38]. In the naïve mouse cerebral cortex, *Ccr5* and *Cxcr2* were among the most broadly expressed receptors across developmental stages, though their expression was not strongly restricted to either Tmem119^+^ or Tmem119^-^ populations.

Overall, four distinct transcriptional programmes, homeostatic, phagocytic, *NRF2/Hmox1* oxidative stress-responsive, and *Apoc1*-associated, were identified and shown to shift in relative dominance across development. Early developmental microglia (E17-P7) were characterized by elevated phagocytic and *NRF2/Hmox1* programme activity, consistent with their roles in synaptic pruning, apoptotic cell clearance, and adaptation to the metabolically demanding environment of the developing brain [2, 3]. With maturation, microglia progressively acquired a homeostatic transcriptional identity marked by *Tmem119* and *P2ry12*, culminating in the stable surveilling phenotype characteristic of the adult brain at P60 [2, 3]. Pseudotime trajectory analysis confirmed this developmental continuum, revealing that the *NRF2/Hmox1* programme peaks at intermediate stages before declining in mature microglia, suggesting a transitional regulatory role for oxidative stress signaling during microglial state transitions. Differential expression analysis further delineated transcriptional signatures distinguishing *Tmem119*^+^ homeostatic microglia from *Tmem119*^-^ populations and identified broad transcriptional remodeling between early developmental and mature microglial states. Additionally, the stage-specific expression of chemokine receptors, particularly *Cxcr4* at early developmental timepoints, points to a role for chemokine-mediated signaling in microglial migration and tissue integration during brain colonization.

A key strength of this study lies in the integration of pseudobulk transcriptomic analysis with hypothesis-driven module scoring, enabling robust characterization of microglial states across development. By focusing on *NRF2/Hmox1*-associated pathways, this work provides new insights into the potential regulatory mechanisms underlying microglial maturation. Future work will extend this analysis to additional developmental stages and integrate single-cell resolution data to further refine microglial state definitions. Ongoing work in our laboratory is actively pursuing these directions, including the incorporation of later post-natal timepoints, inflammatory perturbation datasets, and disease-relevant models, to elucidate how *NRF2/Hmox1*-associated developmental pathways are dysregulated in pathological contexts.

## 6. Conclusion

In conclusion, this study provides a systematic characterization of the apriori defined transcriptional programmes governing microglial development in the naïve mouse cerebral cortex from embryonic day 17 through postnatal day 60. The key homeostatic, phagocytic, *NRF2/Hmox1* oxidative stress response, *Apoc1* associated microglia and chemokine gene expressions in microglia through developmental stages are characterized at naïve state. By integrating PCA, z-scored module scoring, differential expression analysis, pseudotime trajectory modelling, and chemokine receptor profiling, we demonstrate that microglial maturation proceeds along a continuous, coordinated transcriptional axis rather than through discrete, stepwise transitions. Collectively, these findings advance our understanding of the regulatory architecture underlying microglial maturation and highlight the *NRF2/Hmox1* oxidative stress pathway as a developmentally regulated programme with potential relevance to neurological and perinatal brain disorders. This work provides a transcriptional framework that can serve as a reference for future studies investigating how microglial developmental states are perturbed in disease contexts.

## Acknowledgements

We thank Johns Hopkins University (JHU) Department of Pediatrics, Division of Neonatology and JHU Advanced Academic Programs (AAP) Bioinformatics for their continued support.

## Ethical Statement

No animal or human samples were generated for this study.

## Data Accessibility

All data analyzed during this study are included in this published article and its supplementary information files. Raw data is available from National Center for Biotechnology Information (NCBI) BioProject (accession no. PRJNA30727, doi: 10.1073/pnas.1525528113) [6]. Analysis code is provided as supplementary data in this manuscript.

## Authors’ Contributions

Conceptualization and design: VSR, MO, supervision: MO, project administration and funding acquisition: MO, investigation: VSR, MO, methodology: VSR, MO, resources: MO, writing – review and editing: VSR, MO, writing – original draft preparation: VSR, analysis and interpretation: VSR, MO, correspondence and material request: MO. All authors have read and agreed to the published version of the manuscript.

All authors gave final approval for publication and agreed to be held accountable for the work performed therein.

---

**Supplementary Figure 1:**
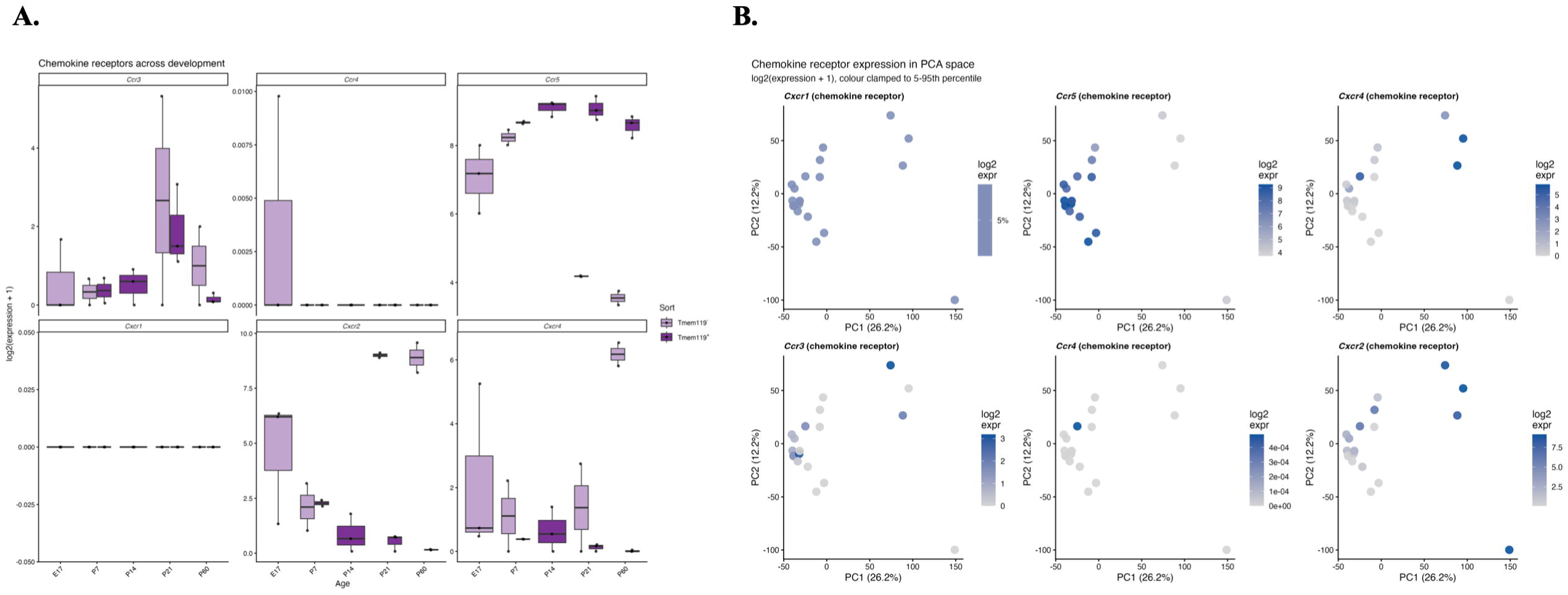
Chemokine Receptor Expression Details. Detailed visualization of chemokine receptor expression (*Cxcr1, Ccr5, Cxcr4, Ccr3, Ccr4, Cxcr2*) across developmental stages using per-gene boxplots and PCA feature plots. **Figure S1A.** Per-gene boxplots of chemokine receptor expression *(Cxcr1, Ccr5, Cxcr4, Ccr3, Ccr4, Cxcr2)* stratified by developmental stage and Tmem119 sort fraction. **Figure S1B.** PCA feature plots for individual chemokine receptors, expression color-clamped to 5th-95th percentile range.

## Supplementary Data

Analysis code is provided within document.

## Supplementary Table

Genome-wide BH-adjusted p-values (FDR): A. DE_tmem119pos_vs_neg(in), B. DE_developmental_vs_mature(in).

